# Size- and colour-based mechanisms shape the phenological structure of butterfly communities

**DOI:** 10.64898/2026.03.26.713911

**Authors:** Roberto Novella-Fernandez, Roland Brandl, Loïc Chalmandrier, Stefan Pinkert, Gerard Talavera, Dirk Zeuss, Christian Hof

## Abstract

1. Seasonal patterns of species appearances constitute a major component of diversity variation. Theory attributes this phenological structuring of communities to the alignment of life cycles to suitable moments and to constraints of seasonality on development, yet the specific mechanisms operating across taxa remain largely unresolved. In insects, body size and colour are key functional traits that contribute to driving spatial community assembly through their link to thermoregulatory performance and development.
2. Here we analyse variation in mean body size and colour lightness of 483 butterfly assemblages across Great Britain and throughout the season to test whether trait alignment with seasonal environment and developmental constraints may shape the phenological structuring of communities.
3. Both body size and body colour varied more along season than across space, emphasizing the importance of phenology on diversity variation. Body size was larger early and late in the season, i.e. under conditions of low temperature and solar radiation. This pattern contrasted with the spatial trends found and was driven by species overwintering as adults, which we interpret as being likely due to energetic constraints. Body colour, conversely, was darker early and late in the season, mirroring the spatial pattern found, and suggesting a thermoregulatory alignment with seasonal conditions. Furthermore, covariation between body size and colour suggests a thermoregulatory interaction between both traits.
4. Our findings suggest that life-cycle constraints and seasonal thermoregulatory alignment contribute to shaping the phenological structure of insect communities.

## Introduction

Many animal groups show diversity variation through the year resulting from species-specific seasonal turnover of life stages. These phenological patterns are thought to arise from seasonal constraints on development (e.g. growth and maturation arrested under cold winter temperatures), together with adaptative alignment of life cycles to seasonal moments of suitable biotic or abiotic conditions that maximise fitness (Chmura et al., 2019; Forrest & Miller-Rushing, 2010; Helm et al., 2013). The mechanisms involved are, however, complex, as such alignment is mediated by species-specific phenological responses to indirect seasonal triggering cues such as photoperiod and temperature (Forrest & Miller-Rushing, 2010; Wolkovich & Donahue, 2021). In addition, developmental constraints and alignment processes interact (Forrest & Miller-Rushing, 2010). As a result, phenological diversity variation remains poorly understood (Shinohara et al., 2023) as compared to spatial diversity variation, which is more straightforwardly driven by filtering processes (abiotic, biotic, dispersal) limiting species’ occurrences (Keddy, 1992). In fact, despite being one of the diversity aspects most directly affected by current global change (Cohen et al., 2018), phenology is often overlooked in animal diversity variation studies (Ponti & Sannolo, 2023), representing a research gap for predicting impacts on biodiversity (Chmura et al., 2019).

Insects are the most diverse animal group. Many major insect groups exhibit pronounced phenological structuring evidenced by a seasonal turnover of adult stages or flight periods (Wolda, 1988). Insect phenological research has largely focused on describing and predicting species-specific development and phenological responses (e.g. adult emergence) to the seasonal cues of temperature and photoperiod (Chuine & Régnière, 2017; Wolkovich & Donahue, 2021). Nevertheless, the drivers behind the phenological structure of insect assemblages remain poorly understood (Chmura et al., 2019, p. 20; Woods et al., 2022). Because species’ traits can drive species’ performance and, thus, fitness under particular environmental conditions, analysing variation in functional trait composition of assemblages across space helps understanding the filtering processes underlying diversity patterns (Tielens & Kelly, 2024). If, as predicted by theory, insect flight periods are adaptatively aligned to seasonal moments of suitable conditions (Helm et al., 2013; Wolkovich & Donahue, 2021), functional composition may also be expected to vary along the season, based on the effect of traits on performance under particular seasonal conditions (Fig1). Analysing functional variation along the season may, thus, help assessing the processes driving phenological structure. This idea was recently supported on dragonfly body colour, which is known to vary spatially in response of thermal conditions, becoming darker in colder areas due to the higher heat gain of dark bodies from solar irradiance (thermal melanism hypothesis; Clusella Trullas et al. 2007, Pinkert et al. 2017). When analysed seasonally, body colour of seasonally-explicit dragonfly assemblages varied aligned with radiation as predicted from thermal melanism (Novella-Fernandez et al., 2023), suggesting that differential colour-based thermoregulation plays a role in driving phenological structure. Together with body colour, size is another thermally relevant trait (Stevenson, 1985) that may be aligned with seasonal thermal conditions. However, observed responses of body size to thermal environment are more complex, as body size of insect assemblages is sometimes larger in colder areas (Bergmann’s Rule; Bergmann 1847, Heinrich 1986), but sometimes it is smaller (Inverse Bergmann’s Rule, e.g. Zeuss et al. 2017). These contrasting patterns may arise because of either the thermal inertial of large bodies, which may explain Bergmann’s rule, or the reduced thermal requirements for flight of small bodies, which may explain the Inverse Bergmann’s rule (Kleckova et al., 2023; May, 1976). However, as body size is also constrained by growth and development (Chown & Gaston, 2010), Bergmann’s pattern may, alternatively, result from warmer temperatures accelerating development more than growth. Consequently, individuals in warm areas may reach the adult stage at smaller sizes (Temperature-Size Rule, TSR; Atkinson 1994, e.g. Chown and Gaston 2010). By contrast, Inverse Bergmann’s rule may result from the shorter favourable season of cold areas which limits the time available for development (Chown & Gaston, 2010). Whether either thermoregulatory performance or developmental constraints on body size may influence the phenological structure of insect assemblages has not been tested.

Butterflies are a diverse and well-known insect taxon that in temperate regions show a replacement of adult flight periods through the season (Scott & Epstein, 1987). Thermal conditions strongly determine butterfly activity, greatly mediated by solar-based thermoregulation (Kingsolver, 1985a), which is in turn facilitated by thermal melanism (Kingsolver, 1985b, 1987, p. 198) as reflected in their colour variation patterns across space (Stelbrink et al., 2019; Zeuss et al., 2014). While synchrony with host plant availability is often considered as a key driver of butterfly phenology (Posledovich et al., 2018), the possible influence of alignment to suitable thermal conditions in determining butterfly flight periods remains unassessed. At the intraspecific level, however, morphological variation along season (seasonal polyphenism) occurs, which has often been suggested to result from alignment to the seasonal environment, e.g in body colouration via thermal melanism (Friberg & Karlsson, 2010; Järvi et al., 2019). Similarly, seasonal polyphenism in body size has been suggested to result from developmental constraints, particularly, the Temperature-Size Rule operating along seasonally changing temperature (Aalberg Haugen et al., 2012; Büyükyilmaz & Tseng, 2022; Friberg & Karlsson, 2010).

Here we analyse trait variation of butterfly assemblages along the season to test whether the trait responses and developmental mechanisms driving community assembly across space may also shape phenological structure. We analyse community-weighted means of body size and colour lightness across sets of 483 assemblages evenly distributed latitudinally across Great Britain and the group’s flight season. First, we assess spatial trait variation. We predict (1A) larger body size with increasing temperature, based on previous predominant support for the Inverse Bergmann’s rule in butterflies (e.g. Barlow 1994, Hawkins and Lawton 1995, Zeuss et al. 2017, Shrestha et al. 2020), and (1B) darker colouration in colder, low-radiation areas, based on previous support for the thermal melanism hypothesis (Stelbrink et al., 2019; Zeuss et al., 2014). We then test whether phenological structure may result from seasonal alignment based on trait-driven thermoregulatory performance, expecting (2A) body size and (2B) colour lightness to vary seasonally in the same direction as their spatial counterparts. Alternatively, if development constrains phenological structure, we expect (3) patterns consistent with the temperature–size rule, with larger summer species whose larvae developed in cool spring conditions, and smaller autumn species whose larvae developed during warm summer conditions.

## Methods

### Building assemblages from occurrence records

We used a curated, expert-validated database of adult butterfly occurrences across Great Britain compiled by the UK Butterfly Monitoring Scheme (UKBMBS https://ukbms.org/). We focused on the period with highest recording intensity 2000 and 2021 and retained ∼ 6.7 million records identified to the species level, with high spatial accuracy (<1000m), and exact dates. From these records, we built 9282 presence/absence assemblages by aggregating observations within ecologically meaningful spatio-temporal scales while controlling for sampling representativeness (Novella-Fernandez et al. 2023, Novella-Fernandez et al. 2024). For that, we bear in mind the definition of assemblage as a set of taxonomically related species co-occurring in space and time and likely to interact (Fauth et al. 1996, Stroud et al. 2015). We first aggregated observations within certain geographical distance (*resSp)*, certain phenological window span (*resPh)*, and certain number of subsequent years (*resTem)*. We then filtered out, from the resulting spatio-phenologically explicit groups of observations, those sufficiently sampled based on a minimum number of sampling events (*samEf)*, a minimum proportion of species recovered based on species accumulation curves (*samCov)*, and minimum species richness (*Smin)*. We evaluated multiple parameter combinations and chose a final set that balanced sample size with fine spatio-temporal resolution: *resSp*=1 km, *resPh*=14 days, *resTem*=0 years, *samEf*=3, *samCov*=0.8, *Smin*=5 species. We consider these as ecologically meaningful assemblages.

To obtain a representative sample of assemblages across latitude and butterflies’ flight season (April-October), we first unclustered assemblages in space and season by keeping a single assemblage within each 10 km grid cell and month. We then grouped assemblages into seven latitudinal groups, spanning from 50.0° to 58.0° N, and kept the minimum number across groups (69). We iterated such unclustering and sampling process to obtain 10 alternative sets of 483 spatio-phenologically independent assemblages that will be used in statistical analyses (Fig. 2a, d). Additionally, to assess the influence of taxonomy and life-cycle strategy on seasonal trait patterns, we built five additional sub-assemblages by excluding, in turn, species of each of the five major taxonomic groups Lycaenidae, Hesperidae, Pieridae, Satyrinae and non-satyrine Nymphalidae (‘Nymphalidae’ for simplicity hereafter). We also built four additional sub-assemblages, each excluding species with one of the four overwintering strategies: egg, larvae, pupa, adult. These sub-assemblages were built using the same parameters as in the overall analysis except for minimum species richness (*Smin),* which was set to 3. The 59 butterfly species included in the assemblages are listed in Table S1. We treated Satyrinae separately from the other Nymphalidae given their deep phylogenetic divergence (Kawahara et al., 2023) and because in our dataset they differ markedly in body size, colouration, and overwintering strategy (Table S2, Fig. S1).

**Figure 1.**
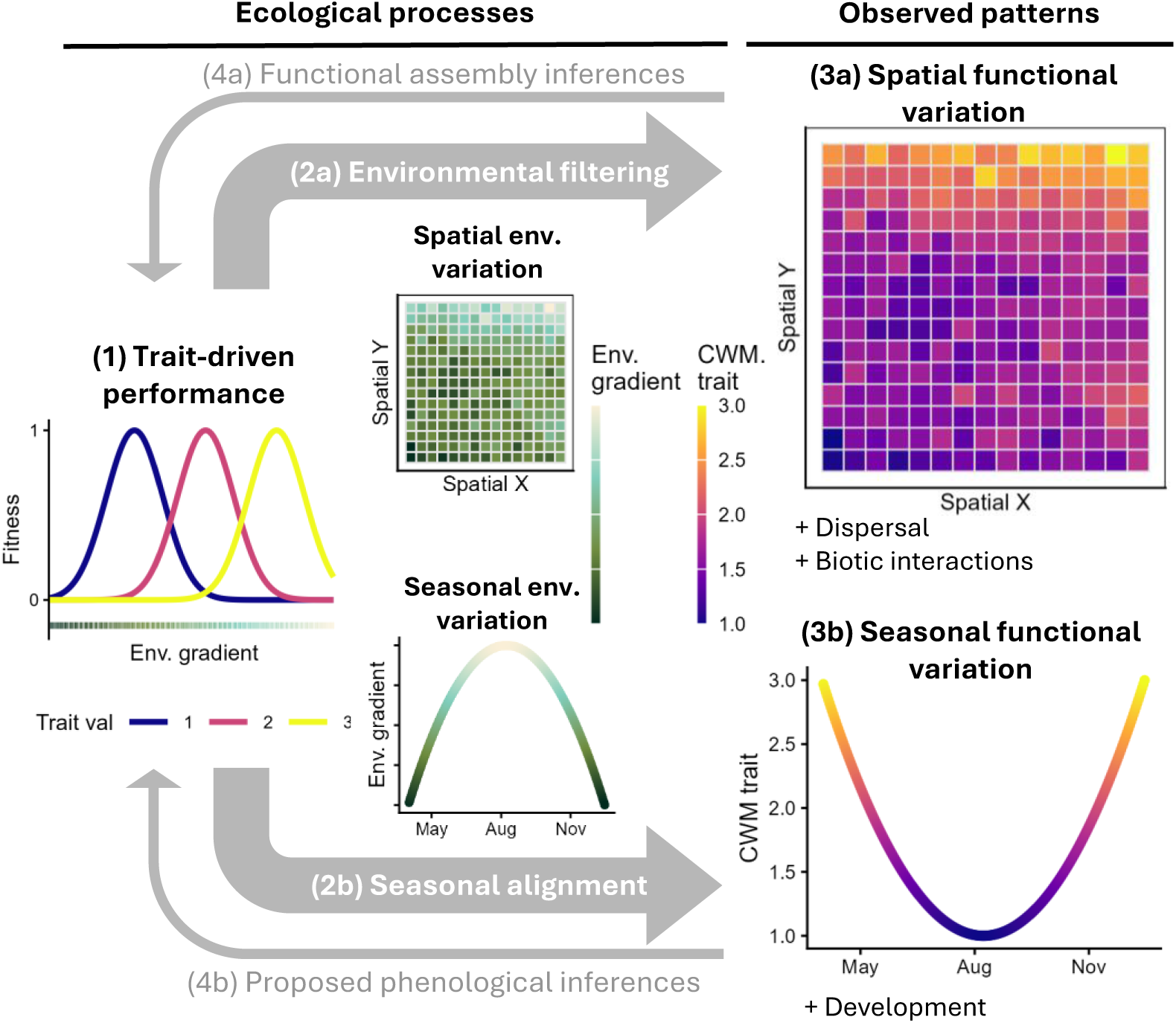
Conceptual framework extending functional inferences from identifying drivers of spatial assembly (a) to identifying drivers of phenological structure (b). Departing from (1) a trait driving differential performance across an environmental gradient (dark green to grey), functional spatial assembly inferences are based on: (2a) Spatial variation in environment filters species based on such trait-driven performance, resulting in (3a) spatial patterns of assemblage-level trait composition, quantified as community-weighted means (CWM), from being dominated by trait value = 1 (dark blue), to being dominated by trait value = 3 (yellow). (4a) Analysing trait variation in relation to the environment can be used to infer environmental filtering. If (2b) life cycles are aligned to seasonal moments of suitable conditions, (3b) variation in traits along the season is also expected based on trait-driven performance. (4b) Analysing such seasonal trait variation in relation to seasonal environmental variation can inform whether an assemblage’s phenological structure results from a trait-driven alignment to suitable seasonal moments. Note that other ecological processes also contribute to shaping both spatial and phenological structures: dispersal limitation and biotic interaction constrain spatial assembly, and development constrains phenological structure.

**Figure 2.**
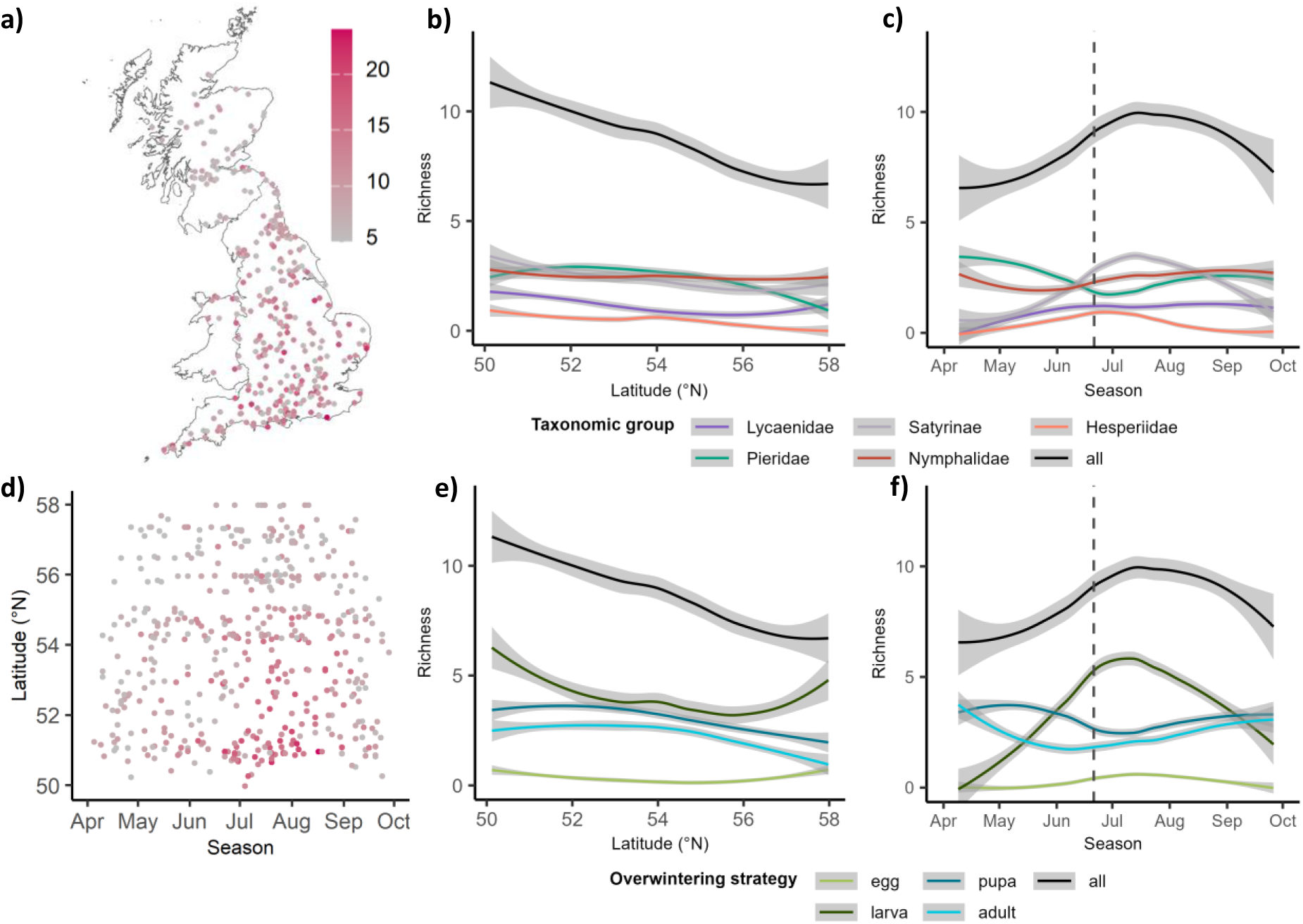
Latitudinal and seasonal distribution of a set of 483 butterfly assemblages and of their species richness. (a) Geographical locations of the studied assemblages across the study area (Great Britain) and (d) their latitudinal distribution across the flight season period. Variation of species richness along latitude (b, e) and season (c, f) depending on taxonomic group (b, c) and overwintering strategy (e, f). The dashed vertical line indicates summer solstice. Only one of the unclustered and stratified sample of assemblages along latitude and season is shown out of the 10 used in statistical analyses (see methods for details).

### Trait data

We quantified butterfly body volume (without wings) and colour lightness using scientific illustrations from Tolman and Lewington (2004) as described by Zeuss *et al*. (2017). Body volume served as a proxy for body size. Colour lightness was estimated from three body regions with expected distinct ecological functions: dorsal area representing mostly wing colouration but including the small proportion of the body (head, thorax and abdomen, <5%), ventral side of the wing area, and body (Stelbrink et al., 2019). These estimates of colour lightness from scientific illustrations were validated to represent an individual’s ability to absorb and reflect radiation (Zeuss et al., 2014). While body size and colour vary mainly among species, they also differ between males and females, despite being significantly correlated (Zeuss et al., 2014). We focused exclusively on females for consistency with previous studies (Stelbrink et al., 2019; Zeuss et al., 2014, 2017). We obtained overwintering stages (egg/larva/pupa/imago) of each species from Willner (2017). Species assigned to multiple categories (9) were included in each of them. We explored variation in species’ trait values across taxonomic groups and overwintering strategies (Fig. S1).

### Replication Statement

| Scale of Inference | Scale at which the factor of interest is applied | Number of replicates at the appropriate scale |
| --- | --- | --- |
| Assemblage | Assemblage | 10 stratified samples of 483 assemblages, distributed across a latitudinal gradient of 8°, and spanning April-October |

### Spatio-seasonal variation in functional assemblage composition along thermal gradients

For each assemblage, we calculated the community weighted means of body size (BS hereafter) and colour lightness of the three body regions: dorsal region (CL_dorsal_), ventral side of wings (CL_ventral_), and body (CL_body_). We analysed variation in all four assemblage-level traits along latitude and across the butterflies’ main flight season period (April-October) by using linear models. Explanatory variables included latitude and day of the year (1-365) (Day), with quadratic effects to better fit non-linear responses.

We obtained raster maps of surface downwelling shortwave radiation and near-surface air temperature at ∼1 km resolution (Karger et al. 2022) for every day of the year between 2005-2014. To characterise the spatio-phenologically explicit conditions of temperature and radiation of assemblages, we first averaged day-to-day temperature and radiation values over the 2005-2014 period and then extracted the corresponding averaged values for each assemblage location and sampling day. To separate the spatial and seasonal components of thermal conditions, we first derived a spatial baseline component for each location. For radiation we averaged values across all days of the year. For temperature we used raster maps of the minimum temperature of the coldest month (BIO6), from WorldClim (https://www.worldclim.org/data/bioclim.html). Then, we derived the seasonal component from the spatio-phenological temperature and radiation values by subtracting their spatial baselines.

To analyse the drivers of assemblages’ functional composition we employed linear models, with BS, CL_dorsal_, CL_ventral_, CL_body_ as response variables depending on the predictor variables spatial temperature and radiation as well as seasonal temperature and radiation. Given that temperature and radiation may influence insect diversity interactively (Bishop et al., 2016), we consider the interactions between spatial temperature and radiation and seasonal temperature and radiation. Additionally, the effects of colour lightness and body size may interact as they both affect thermoregulation (Schweiger & Beierkuhnlein, 2016; Xing et al., 2016) and also covary along season and latitude, thus, they were treated as potentially confounding factors. To assist interpreting their role, we built alternative models for both BS and all three CL response variables, adjusted for the effects of CL traits and BS, respectively. We simplified both raw and adjusted models following a step-wise removal of interactions if they did not increase explanatory power based on AICc. Final raw and adjusted models for BS are shown in Table 1, and for CL traits are shown in Table S3 and Table 2. We assessed the relative contribution of predictors in the variation of each trait using the *lmg* metric from the *relaimpo* R package (Grömping, 2006) which partitions explained variance among predictors. Statistical analyses were replicated across the 10 samples of unclustered and stratified assemblages. Reports on statistical metrics include mean and standard deviation across these 10 replicates. Proportion (0-1) of significant p values (p <0.05) across replicates are given. Significance in less than all replicates (<1) are regarded as non-conclusive trends. Multicollinearity was validated to be within low levels based on variance inflation factor (vif <5). Assumptions of normality of the residuals of statistical models were validated by inspecting diagnostic plots. All variables and thus also the modeĺs β reported are z-standardised to facilitate interpretation.

**Table 1.**
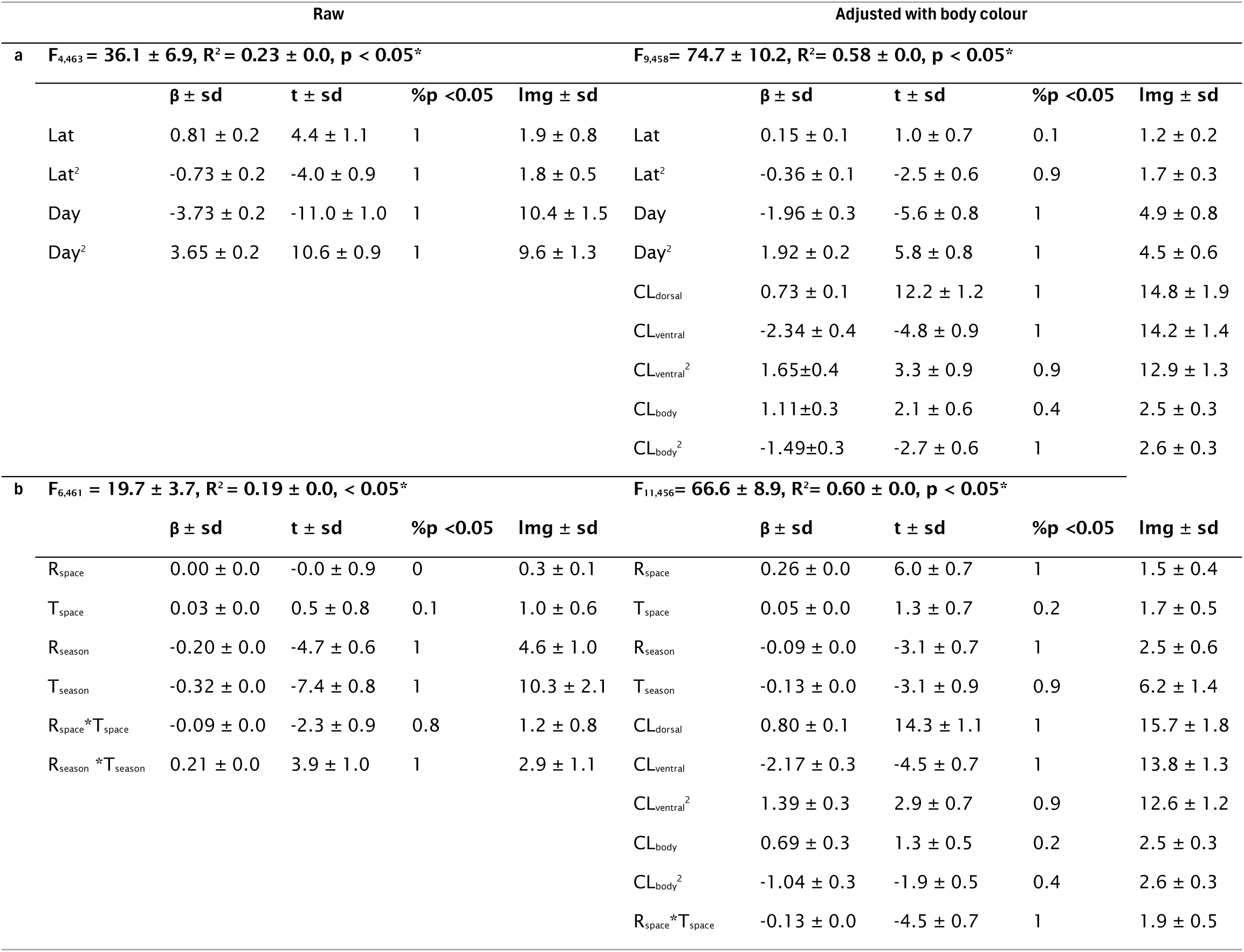
Linear models describing the variation of community weighted means of butterfly body size (BS) along (a) latitude (Lat) and day of the year (Day), as well as in relation to (b) spatio-seasonal thermal variables: spatial temperature (T_space_) and radiation (R_space_), and seasonal temperature (T_season_) and radiation (R_season_). Raw models and adjusted models with the confounding effects of community weighted means of colour lightness (CL_dorsal_: dorsal colour, CL_ventral_: ventral wing colour, CL_body_: body colour) are given. Statistics represent mean ± standard deviation for 10 unclustered and stratified samples of assemblages along latitude and season. %p < 0.05: proportion (0-1) of significant p values (p < 0.05) across those replicates. See methods for detailed description. lmg: Relative importance of the respective predictors as a percentage of variance explained (Grömping, 2006).

| Raw |  |  |  |  | Adjusted with body colour |  |  |  |  |
| --- | --- | --- | --- | --- | --- | --- | --- | --- | --- |
| **a** $F_{4,463} = 36.1 \pm 6.9, R^2 = 0.23 \pm 0.0, p < 0.05^*$ | | | | | $F_{9,458} = 74.7 \pm 10.2, R^2 = 0.58 \pm 0.0, p < 0.05^*$ | | | | |
| | $\beta \pm \text{sd}$ | $t \pm \text{sd}$ | %p < 0.05 | lmg $\pm \text{sd}$ | | $\beta \pm \text{sd}$ | $t \pm \text{sd}$ | %p < 0.05 | lmg $\pm \text{sd}$ |
| Lat | $0.81 \pm 0.2$ | $4.4 \pm 1.1$ | 1 | $1.9 \pm 0.8$ | Lat | $0.15 \pm 0.1$ | $1.0 \pm 0.7$ | 0.1 | $1.2 \pm 0.2$ |
| Lat<sup>2</sup> | $-0.73 \pm 0.2$ | $-4.0 \pm 0.9$ | 1 | $1.8 \pm 0.5$ | Lat<sup>2</sup> | $-0.36 \pm 0.1$ | $-2.5 \pm 0.6$ | 0.9 | $1.7 \pm 0.3$ |
| Day | $-3.73 \pm 0.2$ | $-11.0 \pm 1.0$ | 1 | $10.4 \pm 1.5$ | Day | $-1.96 \pm 0.3$ | $-5.6 \pm 0.8$ | 1 | $4.9 \pm 0.8$ |
| Day<sup>2</sup> | $3.65 \pm 0.2$ | $10.6 \pm 0.9$ | 1 | $9.6 \pm 1.3$ | Day<sup>2</sup> | $1.92 \pm 0.2$ | $5.8 \pm 0.8$ | 1 | $4.5 \pm 0.6$ |
| | | | | | $CL_{\text{dorsal}}$ | $0.73 \pm 0.1$ | $12.2 \pm 1.2$ | 1 | $14.8 \pm 1.9$ |
| | | | | | $CL_{\text{ventral}}$ | $-2.34 \pm 0.4$ | $-4.8 \pm 0.9$ | 1 | $14.2 \pm 1.4$ |
| | | | | | $CL_{\text{ventral}}^2$ | $1.65 \pm 0.4$ | $3.3 \pm 0.9$ | 0.9 | $12.9 \pm 1.3$ |
| | | | | | $CL_{\text{body}}$ | $1.11 \pm 0.3$ | $2.1 \pm 0.6$ | 0.4 | $2.5 \pm 0.3$ |
| | | | | | $CL_{\text{body}}^2$ | $-1.49 \pm 0.3$ | $-2.7 \pm 0.6$ | 1 | $2.6 \pm 0.3$ |
| **b** $F_{6,461} = 19.7 \pm 3.7, R^2 = 0.19 \pm 0.0, < 0.05^*$ | | | | | $F_{11,456} = 66.6 \pm 8.9, R^2 = 0.60 \pm 0.0, p < 0.05^*$ | | | | |
| | $\beta \pm \text{sd}$ | $t \pm \text{sd}$ | %p < 0.05 | lmg $\pm \text{sd}$ | | $\beta \pm \text{sd}$ | $t \pm \text{sd}$ | %p < 0.05 | lmg $\pm \text{sd}$ |
| $R_{\text{space}}$ | $0.00 \pm 0.0$ | $-0.0 \pm 0.9$ | 0 | $0.3 \pm 0.1$ | $R_{\text{space}}$ | $0.26 \pm 0.0$ | $6.0 \pm 0.7$ | 1 | $1.5 \pm 0.4$ |
| $T_{\text{space}}$ | $0.03 \pm 0.0$ | $0.5 \pm 0.8$ | 0.1 | $1.0 \pm 0.6$ | $T_{\text{space}}$ | $0.05 \pm 0.0$ | $1.3 \pm 0.7$ | 0.2 | $1.7 \pm 0.5$ |
| $R_{\text{season}}$ | $-0.20 \pm 0.0$ | $-4.7 \pm 0.6$ | 1 | $4.6 \pm 1.0$ | $R_{\text{season}}$ | $-0.09 \pm 0.0$ | $-3.1 \pm 0.7$ | 1 | $2.5 \pm 0.6$ |
| $T_{\text{season}}$ | $-0.32 \pm 0.0$ | $-7.4 \pm 0.8$ | 1 | $10.3 \pm 2.1$ | $T_{\text{season}}$ | $-0.13 \pm 0.0$ | $-3.1 \pm 0.9$ | 0.9 | $6.2 \pm 1.4$ |
| $R_{\text{space}} * T_{\text{space}}$ | $-0.09 \pm 0.0$ | $-2.3 \pm 0.9$ | 0.8 | $1.2 \pm 0.8$ | $CL_{\text{dorsal}}$ | $0.80 \pm 0.1$ | $14.3 \pm 1.1$ | 1 | $15.7 \pm 1.8$ |
| $R_{\text{season}} * T_{\text{season}}$ | $0.21 \pm 0.0$ | $3.9 \pm 1.0$ | 1 | $2.9 \pm 1.1$ | $CL_{\text{ventral}}$ | $-2.17 \pm 0.3$ | $-4.5 \pm 0.7$ | 1 | $13.8 \pm 1.3$ |
| | | | | | $CL_{\text{ventral}}^2$ | $1.39 \pm 0.3$ | $2.9 \pm 0.7$ | 0.9 | $12.6 \pm 1.2$ |
| | | | | | $CL_{\text{body}}$ | $0.69 \pm 0.3$ | $1.3 \pm 0.5$ | 0.2 | $2.5 \pm 0.3$ |
| | | | | | $CL_{\text{body}}^2$ | $-1.04 \pm 0.3$ | $-1.9 \pm 0.5$ | 0.4 | $2.6 \pm 0.3$ |
| | | | | | $R_{\text{space}} * T_{\text{space}}$ | $-0.13 \pm 0.0$ | $-4.5 \pm 0.7$ | 1 | $1.9 \pm 0.5$ |

**Table 2.**
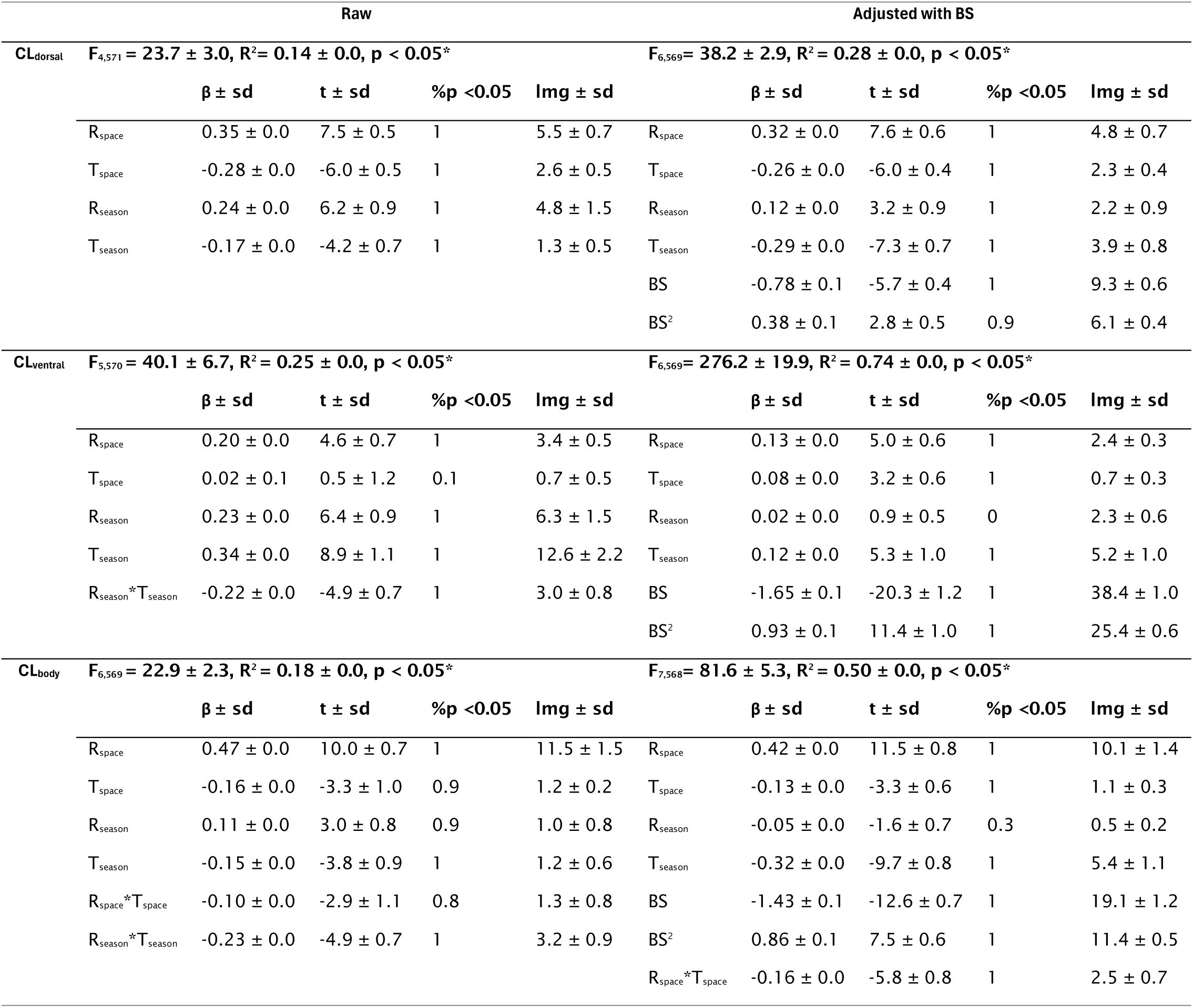
Linear models describing the responses of community weighted means of non-pieridae butterfly assemblages for the three colour lightness traits: CL_dorsal_: dorsal colour, CL_ventral_: ventral wing colour, and CL_body_: body colour, based on spatial temperature (T_space_) and radiation (R_space_), and seasonal temperature (T_season_) and radiation (R_season_). Raw models and adjusted models with the confounding effects of community weighted means of body size (BS). Statistics represent mean ± standard deviation from the 10 unclustered and stratified samples of assemblages along latitude and season. %p <0.05: proportion (0-1) of significant p values (p <0.05) across replicates. See methods for detailed description. lmg: Relative importance of each predictor (Grömping, 2006).

We interpret a negative effect of the spatial components of temperature and radiation on BS as support for Bergmann’s rule, and a positive effect as support of Inverse Bergmann’s rule. Likewise, a positive effect of the spatial components of these variables in CL would support the thermal melanism hypothesis. Similar effects of the seasonal components of the thermal variables on BS or CL to those observed spatially would suggest seasonal alignment based on thermoregulatory performance. Conversely, for BS a pattern of larger body size in summer, whose larvae develop under cold (spring) conditions, and smaller size in autumn, which develop under warm (summer) conditions, would be interpreted as a result from developmental restrictions via the temperature-size rule.

Spatio-phenological patterns may be strongly influenced by specific attributes of certain taxonomic groups or overwintering strategy. Species of the Pieridae family may not respond to thermal melanism as they use reflective thermoregulation (Pieridae: Kingsolver 1985a, b, 1987), and species overwintering as adults may show deviances in body size pattern. To identify the influence of such group effects on the phenological trait variation patterns, we carried out equivalent models of BS and CL in response to thermal drivers separately for all sub-assemblages excluding each of the different taxonomic and overwintering strategy groups. A strong change when excluding a particular group would be interpreted as an influence of such a group-specific contribution to the overall pattern. All analyses were conducted in R.4.4.1 (R Core Team, 2024).

## Results

Our assemblage dataset contained 59 butterfly species: 16 Nymphalidae (excluding Satyrinae), 11 Satyrinae, 14 Lycaenidae, 8 Hesperidae, 8 Pieridae, 1 Papilionidae, and 1 Riodinidae (Table S1). The main overwintering stage strategy was as larva (35), followed by as pupa (14), as egg (11), and finally as adult (7) (Table S2). Body size and the three colour lightness traits differed among taxonomic groups and overwintering strategies. Body size was markedly larger in species that overwinter as adults and in the non-satyrine Nymphalidae (Fig. S1). Colour was consistently lighter in Pieridae across all three body regions, especially in the dorsal region. Conversely, ventral wing colour was darker in species that overwinter as adults and in both Nymphalidae and Satyrinae, with particularly strong differences observed in ventral wing and body coloration (Fig. S1).

Species richness decreased with latitude similarly across taxonomic groups (Fig. 2b) and overwintering strategies (Fig. 2e), with minor deviances from the general trend in some of the groups. Seasonally, richness peaked from late July to early August, but varying across taxonomic groups and overwintering strategies (Fig. 2c, f). In spring, assemblages were dominated by Pieridae and non-satyrine Nymphalidae, while Satyrinae became dominant in summer. By autumn, non-satyrine Nymphalidae and Pieridae regained dominance (Fig. 2c). Finally, Lycaenidae and Hesperidae were never dominant, exhibiting only a weak curvilinear seasonal trend, peaking in mid-summer. Species that overwinter as pupa and adult both dominated in the early and late flight season, whereas larva-overwintering species exhibited a pronounced peak in mid-summer (Fig. 2f).

### Spatio-phenological variation of body size and its drivers

Assemblage-level mean body size (BS) showed minor hump shaped variation with latitude, with largest values at around 54 °N (relative contribution (lmg) Lat + Lat^2^ = 2.7 %, Table 1, Fig. 3a). In contrast, BS showed strong inverse hump-shaped variation along the season, reflecting larger BS in early and late season and smaller BS during summer (lmg Day + Day^2^ = 20 %, Table 1, Fig. 3b). BS was not independent from the mean colour lightness of assemblages (CL) (Table 1). Particularly, larger BS was strongly and non-linearly associated with darker ventral wing colouration (lmg = 27.1 %, Table 1, Fig. S2b), but linearly with lighter dorsal colouration (14.8 %, Table 1, Fig. S2a), as well as mildly and non-linearly with body colouration (5.1 %, Table 1, Fig. S2c). When adjusting the previous model to account for the effects of CL_dorsal_, CL_ventral_, CL_body_, fit improved, with these traits jointly explaining most of BS variation (lmg = 47 %, Table 1), and the variation of BS independent from colour explained by season decreased but remained relevant (lmg Day +Day^2^ = 9.4 %, Table 1). Finally, excluding the 7 species that overwinter as adults from assemblages made seasonal variation in BS to disappear completely (Fig. 2d) but did not change the latitudinal pattern (Fig. 2c).

**Figure 3.**
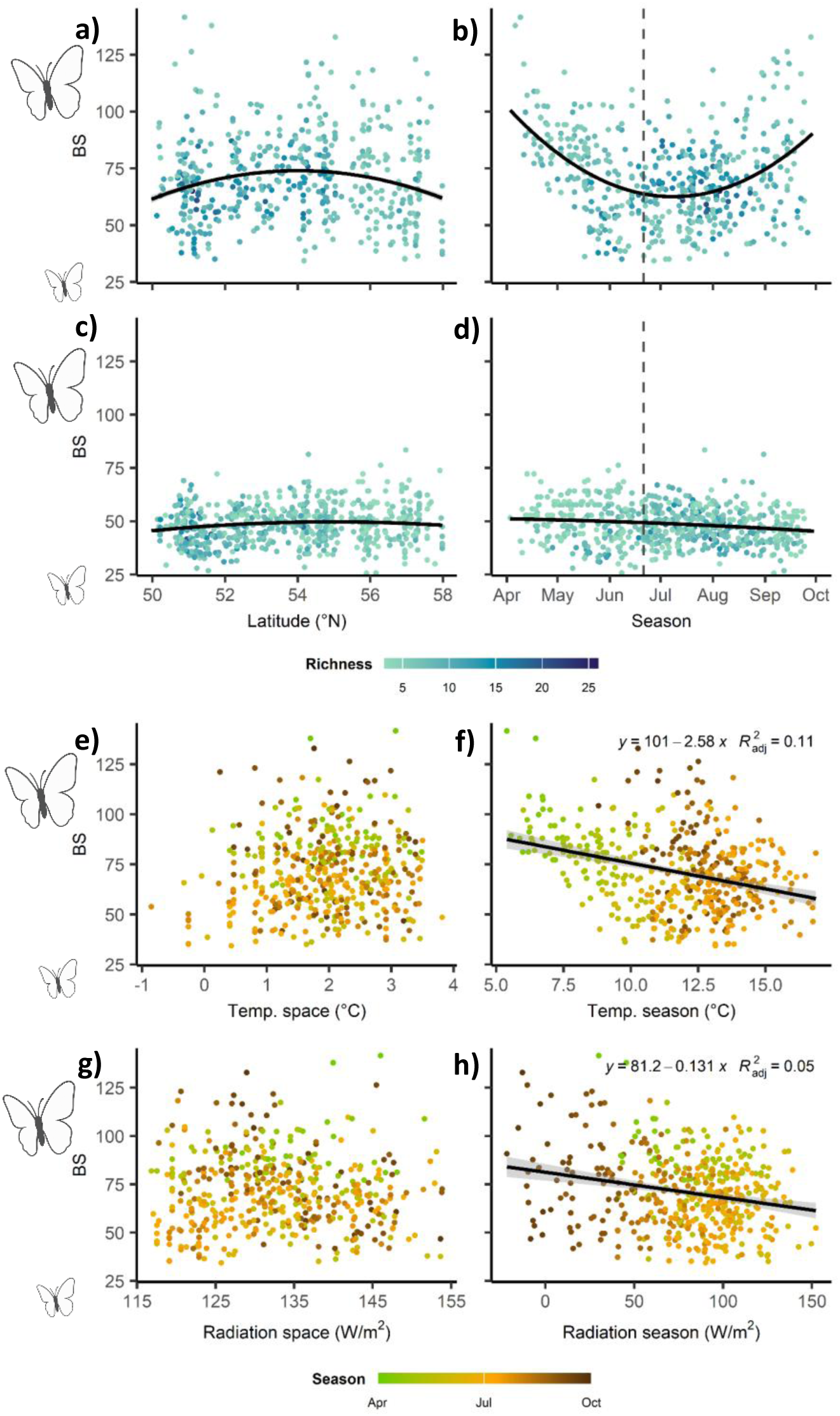
Variation of community weighted means of the body size (BS) of butterfly assemblages. along latitude (a, c) and season (b, d) for the complete assemblages (59 species) (a, b) and excluding the 8 species that overwinter as adults (c, d). Variation of BS in relation to spatial variation of temperature (e) and radiation (g), and seasonal variation of temperature (f) and radiation (h). See methods for details. Points represent a unclustered and stratified samples of assemblages along latitude and season. Black lines in panels a-d describe quadratic relationships across the 10 unclustered and stratified samples of assemblages. Black line in panels e-h represent significant (p < 0.05) relationship of R^2^ > 0.01 in single-variable linear models. Note that significance can differ from multi-variable linear models in Table 1.

BS of the complete assemblages increased marginally in response to the interactive effect of spatial radiation and temperature (joined lmg < 5%, Table 1), yet neither consistently across replicates (80% of replicates p <0.05) nor in single-variable models (Fig. 3e, g). This relationship remained similarly weak, though more complex when adjusting for the confounding effect of the three CL traits (Table 1). In contrast, BS was strongly associated with thermal seasonal conditions (joined lmg = 17.8 %, Table 1), with larger BS associated with lower seasonal temperature (β = -0.32, Fig. 3f) and radiation (β = -0.20, Fig. 3h), with a minor positive interaction between them (Table 1). When adjusting the previous model with the three CL traits, the contribution of seasonal temperature and radiation on BS decreased to moderate levels (joined lmg = 8.7 %).

### Spatio-phenological variation of colour lightness and its drivers

The mean colour of the complete butterfly assemblages was lightest for the ventral side of wings (CL_ventral_: 120.8 **±** 12.2, Fig. 4a), intermediate for the dorsal side (CL_dorsal_: 99.1 **±** 16.9, Fig. 4b), and darkest for the body (CL_body_: 74.6 **±** 8.3, Fig. 4c). All three assemblage-level colouration traits darkened with increasing latitude (Fig. S3a, c, e), more strongly CL_body_ (β = -0.40, R^2^ = 0.16, Fig. S3e) and CL_ventral_ (β = -0.38, R^2^ = 0.14, Fig. S3c), and weaker for CL_dorsal_ (β = -0.20, R^2^ = 0.04, Fig. S3a). Such latitudinal darkening was also apparent when excluding the Pieridae species (Fig. S3b, d, f), which are not strongly reliant on thermal melanism. The spatial component of radiation drove increases in lightness in all three assemblage-level traits (CL_body_: β = 0.61, lmg = 15.0 %. CL_ventral_: β = 0.48, lmg = 12.4 %. CL_dorsal_: β = 0.41, lmg = 5.1 %, Table S3, Fig. S4). Conversely, the spatial component of temperature had only minor contribution (lmg < 2 %) and negative effects (Table S3). Neither adjusting for the effects of BS nor excluding Pieridae did produce large deviations in the effects of spatial components of radiation and temperature in CL traits (Table S3, Table 2, Fig. 5, Fig. S4).

**Figure 4.**
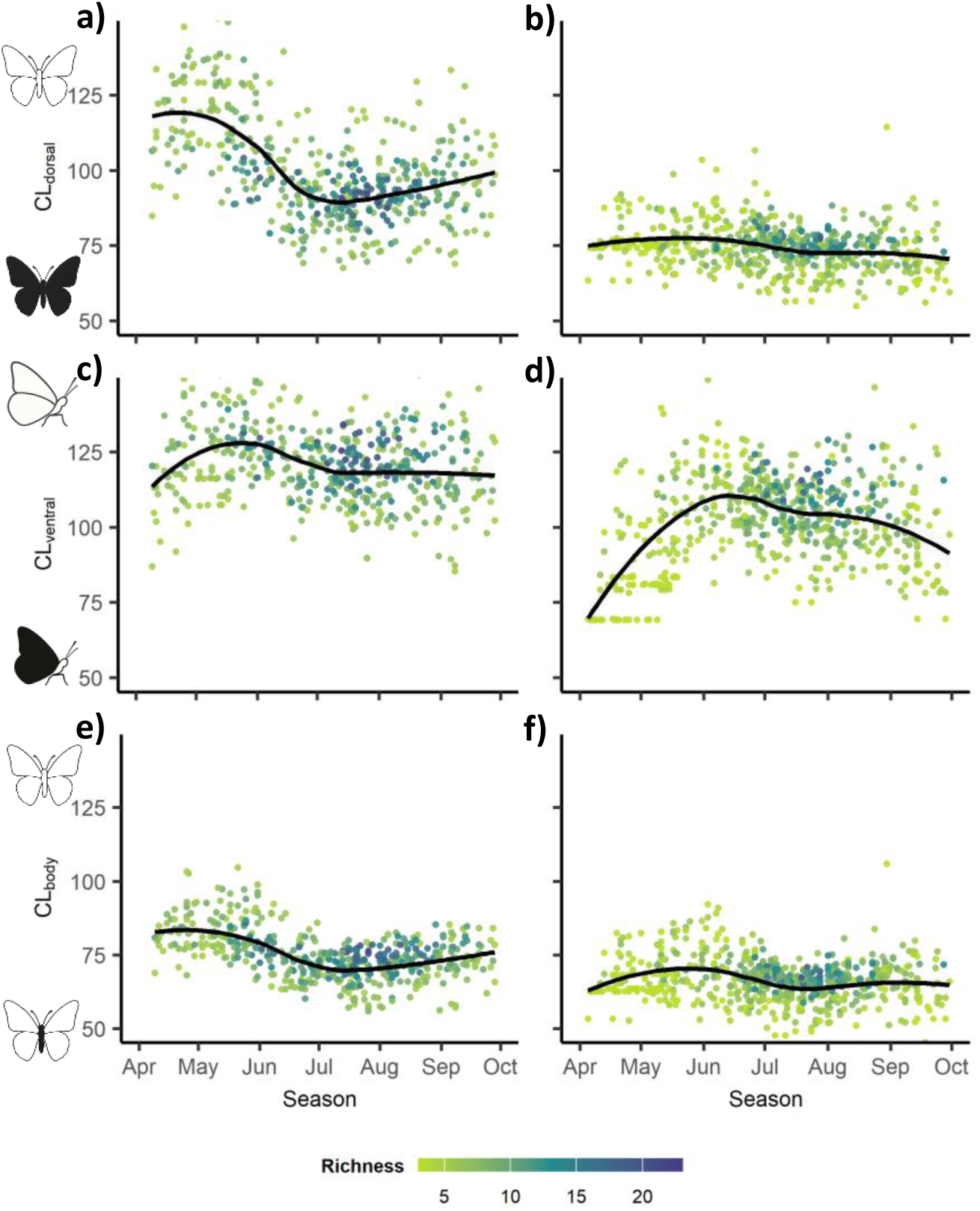
Seasonal variation of community weighted means of the three traits of colour lightness: dorsal side (CL_dorsal_: a, b), ventral wing side (CL_ventral_: c, d), and body (CL_body_: e, f) of complete butterfly assemblages (59 species) (a, c, e), and excluding the 8 Pieridae species, which do not primarily rely on thermal melanism (b, d, f). Points represent a unclustered and stratified sample of assemblages along latitude and season. Black lined describe loess adjustment across the 10 unclustered and stratified samples of assemblages.

**Figure 5.**
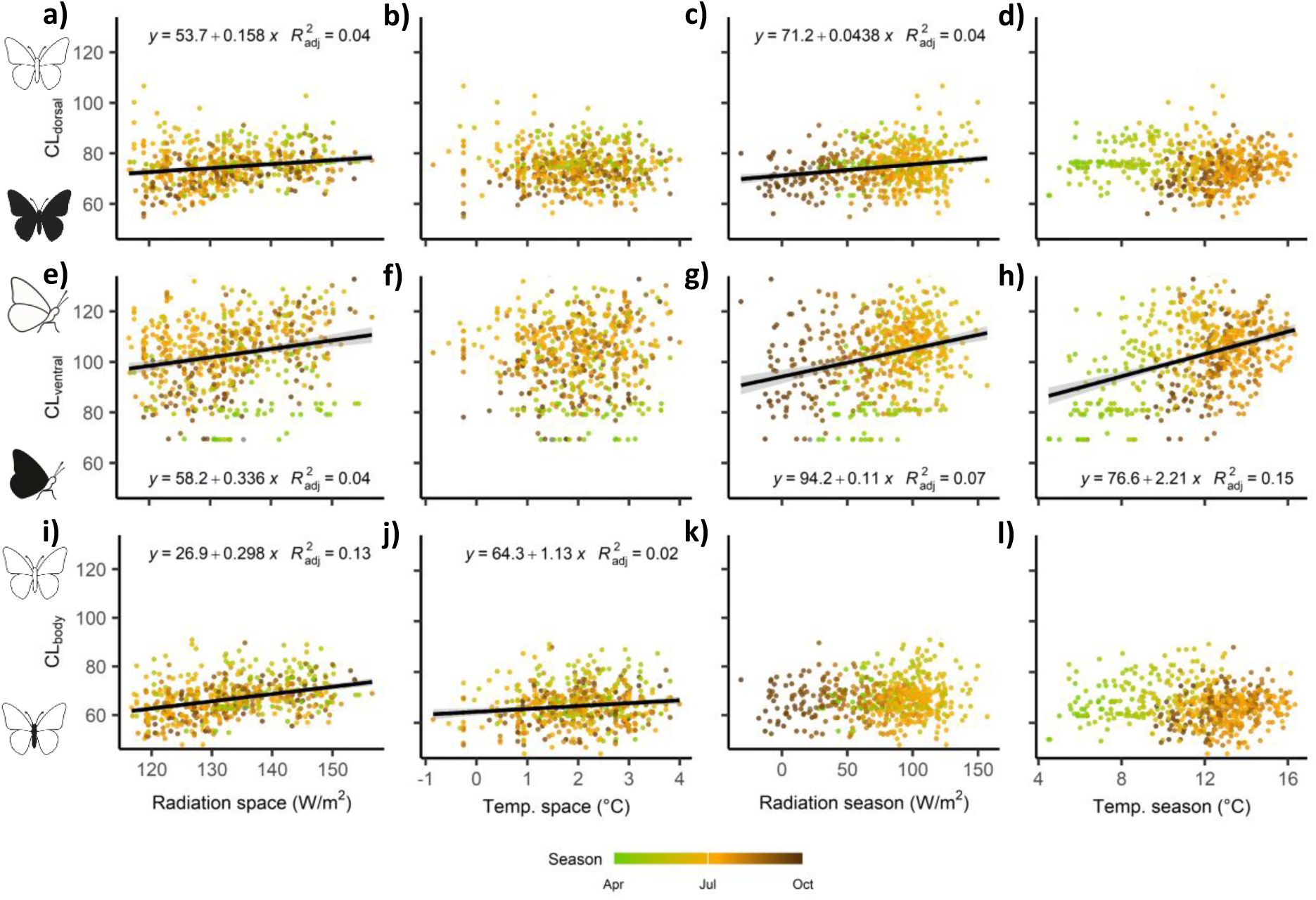
Variation of community weighted means of colouration traits of non-pierid butterfly assemblages. (a-d: CL_dorsal_: dorsal colour, e-h: CL_ventral_: ventral wing colour, i-l: CL_body_: body colour) in relation to the spatial component of radiation (a, e, i) and temperature (b, f, j) and the seasonal component of radiation (c, g, k) and temperature (d, h, l). Points represent an unclustered and stratified sample of assemblages along latitude and season (see methods for more details). Black lines represent significant (p < 0.05) relationships of R^2^ > 0.01 in single-variable linear models. Species of the Pieridae family were excluded as they do not primarily rely on thermal melanism. For responses of the complete butterfly assemblages see Fig. S4.

CL varied largely along the season in all three body regions (Fig. 4), yet strongly differently for the complete assemblages (Fig. 4a, c, e) and when excluding the 8 Pieridae species (Fig. 4b, d, f). In the complete assemblages, CL_dorsal_ followed a strong early-spring peak followed by a sharp decline and moderate lightening from mid-summer (Fig. 4a), CL_ventral_ followed a mild late-spring-peak hump pattern (Fig. 4c), and CL_body_ followed a mild summer-peak inverted hump pattern (Fig. 4e). None of these patterns was aligned with seasonal thermal conditions as predicted from thermal melanism (Table S3). Conversely, seasonal variation in CL_ventral_ of non-Pieridae assemblages was strongly hump-shaped, with lightest colour in June and darkest in early and late seasons (Fig. 4d). In this case, CL_ventra_ was strongly aligned with higher seasonal temperature (β = 0.34, Fig. 5g) and radiation (β = 0.23, Fig. 5h) (joined lmg = 18.9 %, Table 2), with a minor negative interaction (Table 2). Adjusting with the confounding effect of BS reduced the alignment of CL_ventral_ to thermal conditions, preserving only the association with seasonal temperature (β = 0.12, lmg = 5.2 %, Table 2). Seasonal variation of CL_dorsal_ and CL_body_ of non-Pieridae assemblages consisted of a mildly decreasing trend (Fig. 4b, f).

## Discussion

The mechanisms by which insect phenological structures emerge from seasonality remain poorly understood. By analysing seasonal variation in the functional composition of butterfly assemblages, we found pronounced covariation in two essential insect functional traits, body size and colouration that, notably, was stronger than the variation observed across space. These findings highlight phenology as a major, yet often overlooked, component of biodiversity variation. We interpret the observed functional phenological structure within phenological theory as arising from the joint influence of overwintering-related life-cycle constraints and thermoregulatory alignment to seasonal conditions. With this, our study provides insights on the processes through which insect phenological structures arise, opening new avenues for understanding and predicting biodiversity responses to environmental changes.

### Adult overwintering strategy drives seasonal Bergmann’s pattern in butterflies

Mean body size of butterfly assemblages showed little spatial variation, providing unclear support for a spatial Inverse Bergmans’ pattern. This is in line with previous inconsistent spatial body size patterns reported across taxa (Chown & Gaston, 2010). In contrast, body size variation along season was pronounced, with larger-bodied species occurring during seasonal moments of lower radiation and temperature, thus, representing a seasonal expression of Bergmann’s rule. To our knowledge, the only other documented equivalent interspecific pattern is on a wild bee community (Osorio-Canadas et al., 2016). At the intraspecific level, the seasonal Bergmann’s pattern has been more commonly reported e.g. in 88 out of 102 aquatic multivoltine arthropod species exhibited increased body sizes with decreasing temperatures (Horne et al., 2017). While the phenological pattern in Osorio-Canadas et al 2016 was suggested to result from thermoregulatory alignment of body size to season, in our study, the unmatching spatial and seasonal variation of body size in relation to thermal gradients discards that mechanism. In Horne et al. 2017, intra-specific patterns were attributed to developmental constraints based on the temperature-size rule, delaying development in cold seasonal moments. However, because butterfly larval growth is temporarily decoupled from the adult stage, our pattern is not compatible with this neither. Thus, we also discard the temperature size rule as driving seasonal assemblage-level body size variation in butterflies.

Instead, overwintering strategy explained the observed seasonal body size variation, as the pattern disappeared entirely when species overwintering as adults were excluded. Species overwintering as adults – typically larger-bodied and often belonging to the group of non-satyrinae Nymphalidae – were the first to appear in spring and the last in autumn (e.g *Aglais io* and *Aglais urticae*). The relation between large body size and adult overwintering stage has often been found at the intraspecific level, though not universally (Friberg & Karlsson, 2010). For example, the overwintering generation of *Pararge aegeria* has larger body size (Sota 2017), as well as individuals of monarch butterflies found at overwintering sites (Freedman & Dingle, 2018). Adult overwintering butterflies survive the cold winter months remaining inactive in shelters. To endure this period, they pre-accumulate fat reserves (Sinclair and Marshall 2018), which are critical for overwintering survival (Friberg et al. 2023). We interpret that the greater fat-storage capacity of larger-bodied species may facilitate an adult-overwintering life-cycle strategy. Consequently, species overwintering as adults, thus, with late-summer emergence, would have experienced selection favouring larger body sizes to allow sufficient accumulation of energy reserves for winter survival.

### Thermal melanism interacts with body size to drive seasonal variation of butterfly colouration

Butterfly colouration showed a spatial pattern of darker colour in cooler locations, consistent with the thermal melanism hypothesis and in line with our expectations based on larger-scale butterfly studies (Stelbrink et al., 2019; Zeuss et al., 2014). More remarkably, ventral colouration of non-pierid assemblages darkened towards earlier and later seasons coinciding with lower temperature and radiation. These findings coincide with previous results in dragonflies (Novella-Fernandez et al., 2023), and lend support for a seasonal expression of the thermal melanism hypothesis in butterflies, suggesting that differential colour-based thermoregulation may have adaptatively aligned species’ phenologies with thermal conditions, thus, contributing in the phenological structure. Thus, we confirm our expectation that for this trait, the same relationship underpins both spatial and seasonal functional patterns. Despite this assemblage-level pattern is reported here for first time in butterflies, our results, mirror numerous previously reported intraspecific cases of seasonal polyphenism consisting in darker forms later in the season, for instance in *Junonia coenia* (Järvi et al., 2019) *Polygonia c-album* (Nylin, 1992), or *Pararge aegeria* (Van Dyck & Wiklund, 2002), among others (Gautam & Kunte, 2020; Kingsolver & Wiernasz, 1991), which were sometimes suggested to be linked to thermal melanism (Friberg & Karlsson, 2010; Järvi et al., 2019).

The seasonal variation in colouration observed was, however, linked to body size, as assemblages dominated by larger-bodied species also exhibited darker ventral sides of wings, and both assemblage-level traits covaried along the season. This seasonal covariation likely reflects a functional interaction between dark colour and large body size: larger butterflies, with higher wing loading and greater thermal demands, would benefit more from thermal melanism (Heinrich, 1986; Kleckova et al., 2023). Thus, if the species active during the cooler early and late season tend to be larger, their elevated thermal demands would render darker coloration particularly advantageous. Altogether, seasonal variation in body size may largely contribute to the observed seasonal expression of the thermal melanism pattern. A similar association between larger body size and darker colouration was also reported for butterflies in Australia (Xing et al., 2016) and carabid beetles (Schweiger & Beierkuhnlein, 2016). In our study, indeed, the association between larger size and darker colouration became stronger when the Pieridae species, which do not rely strongly on thermal melanism (Kingsolver, 1985b) were excluded.

In addition to thermoregulation, body colour serves many other ecological functions. Improved camouflage during overwintering may also contribute to the colouration in early- and late-season assemblages. Survival during long periods of inactivity for species overwintering as adults may, for instance, be enhanced by darker ventral surfaces that help concealing them from predators. The complete disappearance of the seasonal thermal melanism pattern when including the eight Pieridae species fully meets expectations given the distinct thermoregulatory mechanisms of this group. Pierid butterflies, which are very widespread, fly early in the season and are light-coloured, show an inverse thermal melanism pattern, with a positive relationship between wing lightness and thermal gain, as they reflect radiation towards the body using their pale wings (Kingsolver, 1985b, 1987).

### Limitations

Despite this study focuses on the role of thermoregulatory determinants of phenological structure of insects, we acknowledge that unassessed ecological processes such as biotic ones likely also play essential roles. In butterflies, phenologies of host plants are thought to be important for driving species’ flight periods (Posledovich et al., 2018). Additionally, while the more pronounced functional variation found across season than across space highlights the important contribution of phenology in diversity patterns, a stronger role of space can be expected in study systems spanning larger spatial and thus also abiotic gradients than the relatively narrow variation gradient in Great Britain. Finally, it should be noted that our analyses were constrained by the availability of intraspecific trait data, which limited us to a single trait value per species even though both adult butterfly body size and colouration vary within species. As previously discussed, assemblage-level patterns often mirror intraspecific trends; therefore, incorporating intraspecific functional trait variation may in fact strengthen observed patterns further. Note also, that unaccounted sources of intraspecific variation may influence our patterns. For example, body hair density can reduce heat loss (Kingsolver & Moffat, 1982), or behavioural activity differences across generations (Friberg & Karlsson, 2010; Fric & Konvicka, 2002).

### Conclusions and perspectives in the context of climate change

In the same manner that spatial functional compositional variation can inform of community assembly, by analysing seasonal functional variation within the phenological theorical framework, our study provides insights on the emergence of phenological structure. In particular, we suggest that the phenological structure of butterfly assemblages is adaptatively shaped by the interplay between trait-mediated thermoregulatory alignment with seasonal conditions and developmental constraints. The agreement between our seasonal assemblage-level patterns in both body size and colouration with previously reported intraspecific seasonal trends indicates that the same mechanisms shape phenological patterns across levels of biological organization, echoing other biogeographical patterns, such as for body sizés intraspecific and interspecific spatial variation patterns (Bergmann, 1847; James, 1970). Importantly, our findings have implications for understanding climate change biodiversity impacts, as a thermal melanism seasonal alignment would suggest that butterfly emergence timing is adaptatively mediated to maximise fitness (Bradshaw & Holzapfel, 2007; de Villemereuil et al., 2020) based on thermoregulatory performance. As phenological shifts are among the most widespread climate change responses (Hodgson et al., 2011), such trait-seasonal alignment raises questions regarding the potential for disruption by climate change (Wolkovich & Donahue, 2021). The consequences of such alignment under ongoing and future climate change are, however, beyond our current predictive capacity due to our poor knowledge on the mechanisms and plasticity of phenological cue systems (Chmura et al., 2019; De Lisle et al., 2022). Furthermore, the adaptative potential in both phenological responses and traits themselves is uncertain (Kingsolver & Buckley, 2018). While specific implications for conservation cannot be directly drawn from our study, the suggested importance of thermoregulatory performance on phenological structure stresses the importance of increasing our understanding of the mechanisms behind phenology, phenological alignment across taxa and their adaptive potential via evolution and phenotypic plasticity, particularly in a time of alarming worldwide insect declines (Wagner et al., 2021).

## Acknowledgements

We thank the UKBMS and all contributing observers for their efforts in collecting and providing the observational data. We thank Adrià Miralles for his assistance with butterfly taxonomy. LC acknowledges the support of the ANR-23-IACL-0006 grant.

## Author contributions

RNF: Conceptualisation, Methodology, Software, Formal analysis, Data curation, Writing – original draft, Visualization. RB: Methodology, Writing – review & editing. LC: Methodology, Writing – review & editing. SP: Methodology, Resources, Writing – review & editing. GT: Methodology, Writing – review & editing. DZ: Methodology, Resources, Writing – review & editing. CH: Methodology, Writing – review & editing, Supervision, Funding acquisition

