## Supplementary information for "Size- and colour-based mechanisms shape the phenological structure of butterfly communities"

For

### TABLES

**Table S1.** List of the 59 butterfly species included in the study.

| Species | Family/Subfamily | Species | Family/Subfamily |
|---|---|---|---|
| *Carterocephalus palaemon* | Hesperiidae | *Thecla betulae* | Lycaenidae |
| *Erynnis tages* | Hesperiidae | *Hamearis lucina* | Riodinidae |
| *Hesperia comma* | Hesperiidae | *Aglais io* | Nymphalidae (Nymphalinae) |
| *Pyrgus malvae* | Hesperiidae | *Boloria (Clossiana) euphrosyne* | Nymphalidae (Heliconiinae) |
| *Thymelicus acteon* | Hesperiidae | *Boloria (Clossiana) selene* | Nymphalidae (Heliconiinae) |
| *Thymelicus lineola* | Hesperiidae | *Euphydryas aurinia* | Nymphalidae (Nymphalinae) |
| *Thymelicus sylvestris* | Hesperiidae | *Fabriciana adippe* | Nymphalidae (Heliconiinae) |
| *Ochlodes venata* | Hesperiidae | *Melitaea athalia* | Nymphalidae (Nymphalinae) |
| *Papilio machaon* | Papilionidae | *Nymphalis polychloros* | Nymphalidae (Nymphalinae) |
| *Pieris napi* | Pieridae | *Speyeria aglaja* | Nymphalidae (Heliconiinae) |
| *Pieris rapae* | Pieridae | *Aglais urticae* | Nymphalidae (Nymphalinae) |
| *Anthocharis cardamines* | Pieridae | *Apatura iris* | Nymphalidae (Apaturinae) |
| *Colias croceus* | Pieridae | *Argynnis paphia* | Nymphalidae (Heliconiinae) |
| *Colias hyale* | Pieridae | *Limenitis camilla* | Nymphalidae (Limenitidinae) |
| *Gonepteryx rhamni* | Pieridae | *Melitaea cinxia* | Nymphalidae (Nymphalinae) |
| *Leptidea sinapis* | Pieridae | *Polygonia c-album* | Nymphalidae (Nymphalinae) |
| *Pieris brassicae* | Pieridae | *Vanessa atalanta* | Nymphalidae (Nymphalinae) |
| *Quercusia quercus* | Lycaenidae | *Vanessa cardui* | Nymphalidae (Nymphalinae) |
| *Aricia agestis* | Lycaenidae | *Aphantopus hyperantus* | Nymphalidae (Satyrinae) |
| *Aricia artaxerxes* | Lycaenidae | *Coenonympha pamphilus* | Nymphalidae (Satyrinae) |
| *Callophrys rubi* | Lycaenidae | *Coenonympha tullia* | Nymphalidae (Satyrinae) |
| *Celastrina argiolus* | Lycaenidae | *Erebia aethiops* | Nymphalidae (Satyrinae) |
| *Cupido minimus* | Lycaenidae | *Erebia epiphron* | Nymphalidae (Satyrinae) |
| *Lycaena phlaeas* | Lycaenidae | *Hipparchia semele* | Nymphalidae (Satyrinae) |
| *Lysandra bellargus* | Lycaenidae | *Lasiommata megera* | Nymphalidae (Satyrinae) |
| *Lysandra coridon* | Lycaenidae | *Maniola jurtina* | Nymphalidae (Satyrinae) |
| *Plebejus argus* | Lycaenidae | *Melanargia galathea* | Nymphalidae (Satyrinae) |
| *Polyommatus icarus* | Lycaenidae | *Pararge aegeria* | Nymphalidae (Satyrinae) |
| *Fixsenia pruni* | Lycaenidae | *Pyronia tithonus* | Nymphalidae (Satyrinae) |
| *Satyrium w-album* | Lycaenidae | | |

**Table S2.** Number of species per overwintering strategy across taxonomic groups.

| | Egg | Larva | Pupa | Adult |
|---|---|---|---|---|
| Hesperiidae | 2 | 6 | 1 | 0 |
| Lycaenidae | 6 | 6 | 2 | 0 |
| Non-Satyrinae Nymphalidae | 3 | 10 | 1 | 6 |
| Papilionidae | 0 | 0 | 1 | 0 |
| Pieridae | 0 | 2 | 7 | 1 |
| Riodinidae | 0 | 0 | 1 | 0 |
| Satyrinae | 0 | 11 | 1 | 0 |

**Table S3.** Linear model describing the responses of community weighted means of the complete butterfly assemblages for the three colour lightness traits: $CL_{dorsal}$: dorsal colour, $CL_{ventral}$: ventral wing colour, and $CL_{body}$: body colour, based on spatial temperature ($T_{space}$) and radiation ($R_{space}$), and seasonal temperature ($T_{season}$) and radiation ($R_{season}$). Raw models and adjusted models with the confounding effects of community weighted means of body size (BS). Statistics represent mean ± standard deviation from the 10 unclustered and stratified samples of assemblages along latitude and season. %p<0.05: proportion (0-1) of significant p values (p <0.05) across replicates. See methods for detailed description. lmg: Relative importance of each predictor (Grömping 2006).

| | Raw | | | | | Adjusted with BS | | | | |
|---|---|---|---|---|---|---|---|---|---|---|
| $CL_{dorsal}$ | $F_{6,461}$= 50.4 ± 8.5, $R^2$= 0.39 ± 0.0, p<0.05* | | | | | | | | | |
| | | β ± sd | t ± sd | %p <0.05 | lmg ± sd | | | | | |
| | $R_{space}$ | 0.41 ± 0.0 | 9.1 ± 1.3 | 1 | 5.1 ± 1.8 | | | | | |
| | $T_{space}$ | -0.30 ± 0.0 | -6.6 ± 1.1 | 1 | 1.6 ± 0.6 | | | | | |
| | $R_{season}$ | 0.10 ± 0.0 | 2.6 ± 0.5 | 0.8 | 0.4 ± 0.2 | | | | | |
| | $T_{season}$ | -0.59 ± 0.0 | -15.6 ± 1.2 | 1 | 28.6 ± 2.5 | | | | | |
| | $R_{space}*T_{space}$ | -0.14 ± 0.0 | -4.2 ± 0.5 | 1 | 2.1 ± 0.5 | | | | | |
| | $R_{season}*T_{season}$ | -0.16 ± 0.0 | -3.6 ± 0.8 | 1 | 1.5 ± 0.6 | | | | | |
| $CL_{ventral}$ | $F_{6,461}$= 22.8 ± 4.8, $R^2$= 0.22 ± 0.0, p <0.05* | | | | | $F_{7,460}$= 52.2 ± 9.3, $R^2$= 0.43 ± 0.0, p<0.05* | | | | |
| | | β ± sd | t ± sd | %p <0.05 | lmg ± sd | | β ± sd | t ± sd | %p <0.05 | lmg ± sd |
| | $R_{space}$ | 0.48 ± 0.1 | 9.4 ± 1.4 | 1 | 12.4 ± 3.4 | $R_{space}$ | 0.48 ± 0.0 | 11.1 ± 1.3 | 1 | 12.2 ± 2.9 |
| | $T_{space}$ | -0.14 ± 0.0 | -2.6 ± 0.8 | 0.8 | 1.3 ± 0.3 | $T_{space}$ | -0.12 ± 0.0 | -2.8 ± 0.8 | 0.9 | 1.4 ± 0.3 |
| | $R_{season}$ | 0.17 ± 0.0 | 4.2 ± 0.8 | 1 | 2.1 ± 0.7 | $R_{season}$ | 0.07 ± 0.0 | 2.0 ± 0.6 | 0.5 | 1.3 ± 0.5 |
| | $T_{season}$ | -0.14 ± 0.0 | -3.2 ± 1.0 | 0.9 | 1.3 ± 0.9 | $T_{season}$ | -0.30 ± 0.0 | -7.9 ± 0.9 | 1 | 3.3 ± 1.0 |
| | $R_{space}*T_{space}$ | -0.11 ± 0.0 | -2.9 ± 0.7 | 0.9 | 1.3 ± 0.6 | BS | -0.51 ± 0.0 | -13.2 ± 1.5 | 1 | 21.1 ± 3.7 |
| | $R_{season}*T_{season}$ | -0.27 ± 0.0 | -5.3 ± 0.5 | 1 | 4.4 ± 0.9 | $R_{space}*T_{space}$ | -0.16 ± 0.0 | -4.8 ± 0.5 | 1 | 2.4 ± 0.6 |
| | | | | | | $R_{season}*T_{season}$ | -0.17 ± 0.0 | -3.7 ± 0.5 | 1 | 2.2 ± 0.4 |
| $CL_{body}$ | $F_{6,461}$= 63.8 ± 10.3, $R^2$= 0.44 ± 0.0, p <0.05* | | | | | $F_{7,460}$= 69.8 ± 11.1, $R^2$= 0.50 ± 0.0, p < 0.05* | | | | |
| | | β ± sd | t ± sd | %p <0.05 | lmg ± sd | | β ± sd | t ± sd | %p <0.05 | lmg ± sd |
| | $R_{space}$ | 0.61 ± 0.0 | 14.1 ± 1.5 | 1 | 15.0 ± 3.0 | $R_{space}$ | 0.61 ± 0.0 | 14.9 ± 1.5 | 1 | 14.9 ± 2.9 |
| | $T_{space}$ | -0.31 ± 0.0 | -7.1 ± 1.1 | 1 | 2.0 ± 0.3 | $T_{space}$ | -0.30 ± 0.0 | -7.3 ± 1.1 | 1 | 1.9 ± 0.3 |
| | $R_{season}$ | 0.09 ± 0.0 | 2.5 ± 0.7 | 0.8 | 0.3 ± 0.1 | $R_{season}$ | 0.03 ± 0.0 | 1.0 ± 0.6 | 0 | 0.2 ± 0.1 |
| | $T_{season}$ | -0.53 ± 0.0 | -14.7 ± 1.4 | 1 | 21.7 ± 2.5 | $T_{season}$ | -0.62 ± 0.0 | -17.2 ± 1.4 | 1 | 23.8 ± 2.4 |
| | $R_{space}*T_{space}$ | -0.20 ± 0.0 | -6.2 ± 0.6 | 1 | 4.3 ± 0.8 | BS | -0.27 ± 0.1 | -7.5 ± 1.5 | 1 | 4.2 ± 1.7 |
| | $R_{season}*T_{season}$ | -0.19 ± 0.0 | -4.3 ± 0.5 | 1 | 1.9 ± 0.4 | $R_{space}*T_{space}$ | -0.22 ± 0.0 | -7.3 ± 0.5 | 1 | 5.1 ± 0.7 |
| | | | | | | $R_{season}*T_{season}$ | -0.13 ± 0.0 | -3.2 ± 0.5 | 1 | 1.1 ± 0.3 |

### FIGURES

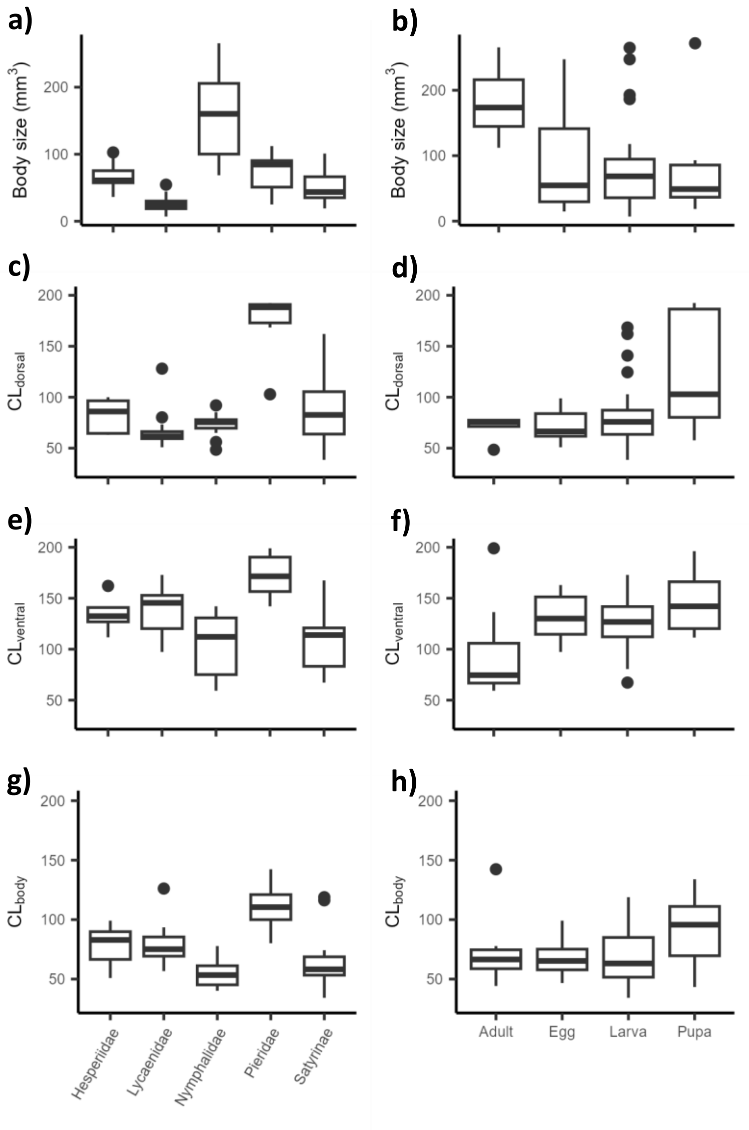


**Figure S1.** Trait values of species body size (BS) and colour lightness of dorsal side ($CL_{dorsal}$), ventral side of wings ($CL_{ventral}$), and body ($CL_{body}$) across taxonomic groups and overwintering strategies. Note that the Nymphalidae group excludes the Satyrinae.

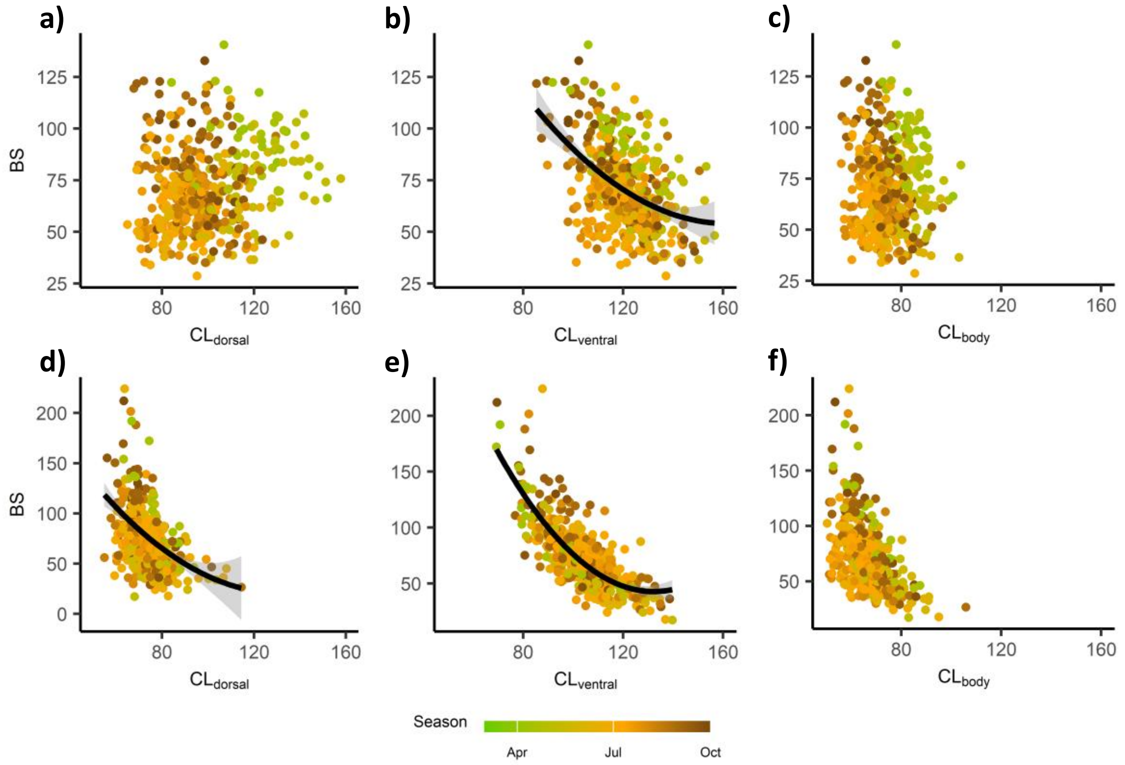


**Figure S2.** Relation between community weighted means of body size (BS) and colour lightness of dorsal side ($CL_{dorsal}$), ventral side of wings ($CL_{ventral}$), and body ($CL_{body}$) for butterfly assemblages. Results shown in the top row (a-c) are based on all 59 species and families of Great Britain and those in the bottom row (d-f) are based on assemblages excluding all 8 species of the family Pieridae. Note that significance can differ from multi-variable linear models in Table 1 and Table 2.

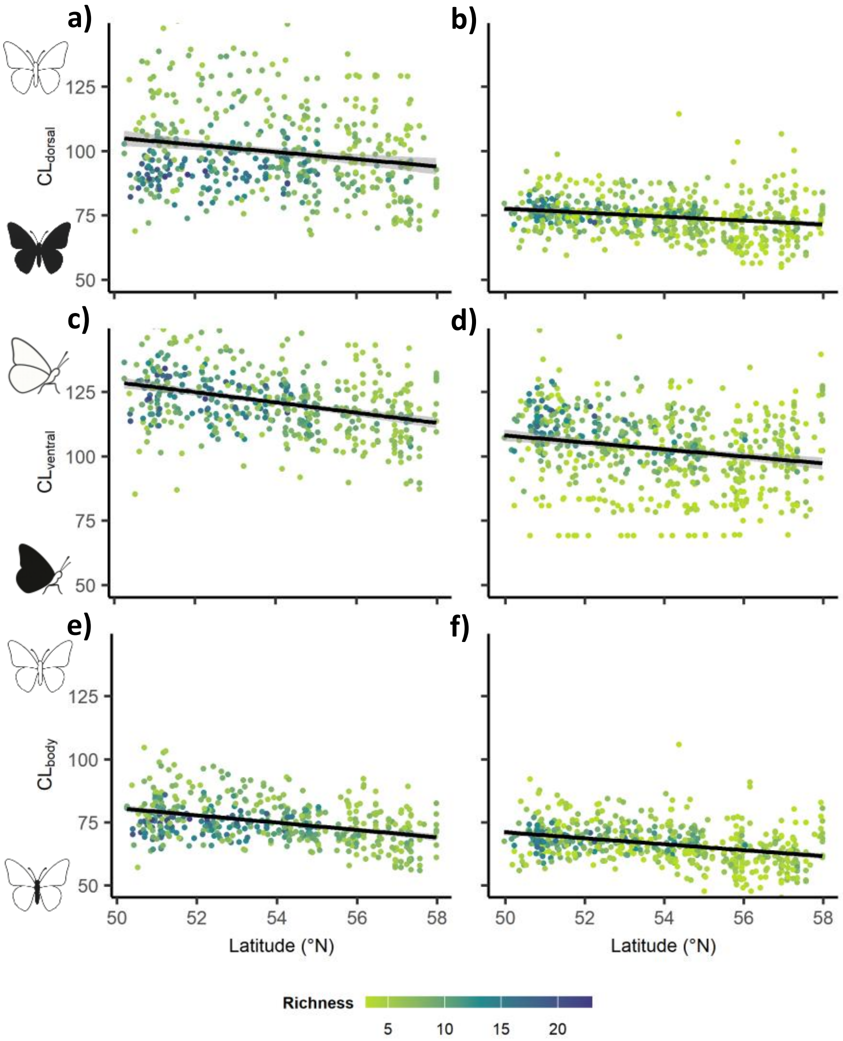


**Figure S3.** Latitudinal variation in colour ligthness of butterfly assemblages colour lightness of dorsal side ($CL_{dorsal}$), ventral side of wings ($CL_{ventral}$), and body ($CL_{body}$) along season for the complete butterfly assemblages (a, c, e) and excluding the Pieridae (b, d, f). Black lines represent linear models: a) $F_{1,466} = 20.4 \pm 10.5$, $R^2 = 0.04 \pm 0.0$, $\beta = -0.20 \pm 0.0$, $t = -4.4 \pm 1.1$, % $p < 0.05 = 1$. b) $F_{1,574} = 27.8 \pm 5.2$, $R^2 = 0.04 \pm 0.0$, $\beta = -0.21 \pm 0.0$, $t = -5.3 \pm 0.5$, % $p < 0.05 = 1$. c) $F_{1,466} = 78.8 \pm 22.8$, $R^2 = 0.14 \pm 0.0$, $\beta = -0.38 \pm 0.0$, $t = -8.8 \pm 1.3$, % $p < 0.05 = 1$. d) $F_{1,574} = 22.5 \pm 4.1$, $R^2 = 0.03 \pm 0.0$, $\beta = -0.19 \pm 0.0$, $t = -4.7 \pm 0.4$, % $p < 0.05 = 1$. e) $F_{1,466} = 89.4 \pm 24.8$, $R^2 = 0.16 \pm 0.0$, $\beta = -0.40 \pm 0.0$, $t = -9.4 \pm 1.3$, % $p < 0.05 = 1$. f) $F_{1,574} = 83.6 \pm 13.3$, $R^2 = 0.12 \pm 0.0$, $\beta = -0.36 \pm 0.0$, $t = -9.1 \pm 0.7$, % $p < 0.05 = 1$.

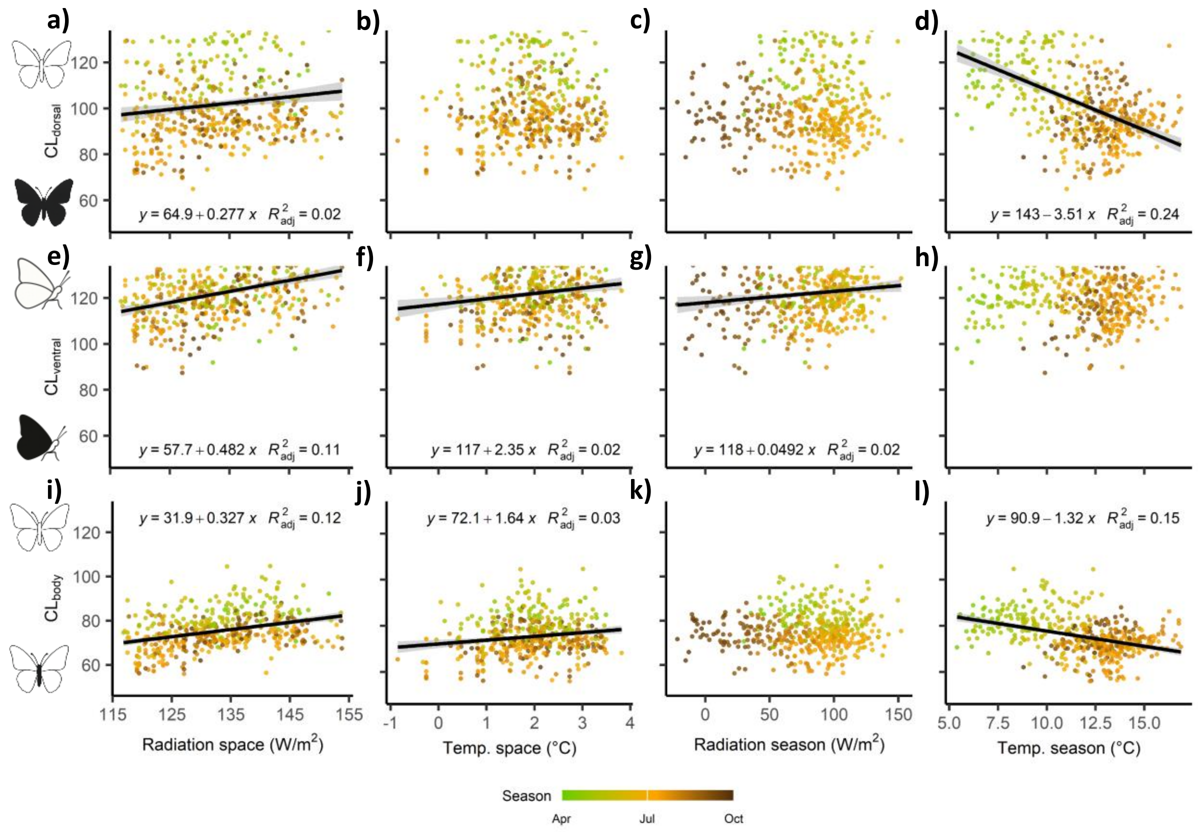


**Figure S4.** Variation of community weighted means of colouration traits of the complete butterfly assemblages (a-d: $CL_{dorsal}$: dorsal colour, e-h: $CL_{ventral}$: ventral wing colour, i-l: $CL_{body}$: body colour) in relation to the spatial component of radiation (a, e, i) and temperature (b, f, j) and in relation to the seasonal component of radiation (c, g, k) and temperature (d, h, l). Points represent an unclustered and stratified sample of assemblages along latitude and season. See methods for more details. Black lines represent significant ($p < 0.05$) relationship with $R^2 > 0.01$ in single-variable linear models.
